# Blood Chemistry Analysis in an Active Pneumocystis Pneumonia (PCP) Model Treated with Selective CARD9 Inhibitor BRD5529

**DOI:** 10.1101/2022.07.21.500972

**Authors:** Theodore J. Kottom, Kyle Schaefbauer, Andrew H. Limper

## Abstract

**Background:** Pneumocystis pneumonia (PCP) in AIDS and other immunosuppressive states that result from absence of CD4 lymphocytic immunity, continues to be a significant cause of morbidity and mortality. We and others have shown the importance of CARD9 in PCP and other fungal infections, respectively. BRD5529 has been shown to be an effective in vitro and in vivo (18 hour) inhibitor of *Pneumocystis* β-glucans induced proinflammatory response. These recent results, along with recent general safety and toxicology assessments suggests the application of BRD5529 in an active PCP mouse model of infection to assess initial blood toxicology parameters.

**Methods:** To assess preliminary blood toxicology, mice were injected intraperitoneally (IP) daily either with vehicle or BRD5529 at 1.0 mg/kg for one week starting at the 6th week of the PCP mouse model. After one week, mice were sacrificed, and blood collection postmortem was performed for blood chemistry analysis.

**Results:** Analysis of blood chemistry showed a significant reduction in blood urea nitrogen (BUN) in the BRD5529 IP treated PCP mice cohort compared to the vehicle control group. All other blood chemistry parameters were not significantly different between the two groups.

**Conclusions:** BRD5529 in this preliminary PCP treatment model displayed only significant changes in BUN levels in the BRD5529 treatment group versus the vehicle control group. All other blood chemistry parameters were statistically similar between the two groups.

## 1. Introduction

We have previously shown that host innate immunity to *Pneumocystis* is mediated by C-Type Lectin Receptors (CLRs) on macrophages and involves downstream CARD9 activation^1^. We have further shown that the CARD9 downstream initiated proinflammatory response via *Pneumocystis* β-glucans can be targeted by a novel specific small molecule inhibitor of CARD9 termed BR5529 both in vitro and in vivo^2,3^. The purpose of this study was to evaluate if treatment of a 6-week PCP mouse model with BRD5529 resulted in significant changes in blood chemistry analysis as compared to the similar infected cohort vehicle control group.

## 2. Methods

### 2.1 Animals

Equal number of male and female C57BL/6 mice (The Jackson Laboratory) at 10-12 weeks of age were used in this study. Animal procedures were performed according to the Laboratory Animal Welfare Act, and the Mayo Clinic Institutional Animal Care and Use Committee (IACUC) (Approval number: A00005722020).

### 2.2 BRD5529 administration

Methocel™ and BRD5529 was obtained from Sigma Aldrich. Methocel™ was used to prepare BRD5529 due to the compounds lack of solubility in water or saline^4^. The inhibitor was prepared with 1% Methocel™ ^2^. Intraperitoneal treatment (100 ul) with 1% Methocel™ (vehicle, control mice group) or 1 mg/kg of BRD5529 inhibitor in Methocel™ was initiated on day 42 of an active PCP mouse model infection. IP injections were given daily for 7 days. At day 49 of the PCP infection, mice were sacrificed, and blood chemistry analysis conducted as described below.

### 2.3 Blood chemistry analysis

Analysis of blood chemistry was conducted with the Piccolo Xpress™ Chemistry Analyzer according to the manufacturer’s instructions.

### 2.4 Statistical analysis

2-sample unpaired Student’s *t*-test was utilized. Analysis of data was conducted on Prism 9 for macOS, version 9.4.0 (GraphPad). Values of *p* < 0.05 were considered significant.

## 3. Results

### 3.1 Serum chemistry data

Two group mean serum chemistry data from day 49 of PCP with treatment of animals with either 1mg/kg BRD5529 or vehicle control daily for 7 days starting at day 42 of the PCP model. Of the 13 blood parameters tested, only BUN levels were noted significantly lower in the vehicle control treated group compared to the BRD5529 group (Fig. 1 A-K).

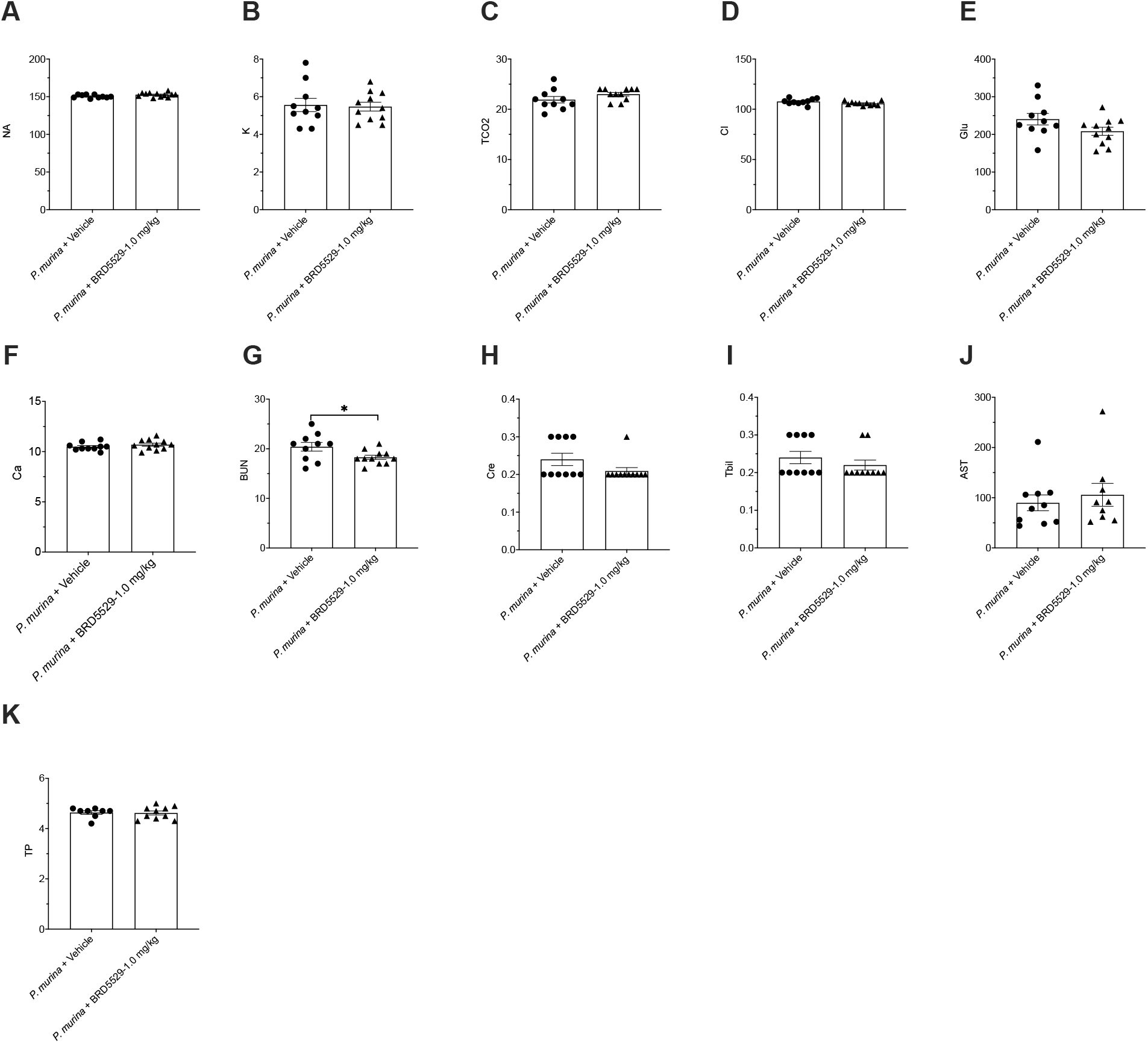
Na, sodium (mmol/L); K, potassium (mmol/L); TCO_2_, carbon dioxide (mmol/L); Cl, chloride (mmol/L); Glu, glucose (mg/dL); Ca, Calcium (mg/dL); BUN, blood urea nitrogen (mg/dL); Cre, creatine (g/dL); Tbil, total bilirubin (mg/dL); AST, aspartate aminotransferase (U/L); TP, total protein (g/dL). (n = 10-11 mice/group). *p<0.05

## 4. Discussion

Trimethoprim-sulfamethoxazole (TMP-SMX) is the most common anti-*Pneumocystis* therapy, and the combination has shown great effectiveness in treating *Pneumocystis jirovecii* pneumonia (PJP). Unfortunately, upon organism killing with antifungal therapy, highly proinflammatory β- glucan carbohydrate is also exposed and released from the cyst cell wall, resulting in highly detrimental results to the host ^5-7^. Therefore, other adjunct therapies might be considered in PJP treatment. We have recently published that the caspase recruitment domain-containing protein 9 (CARD9) inhibitor BRD5529 can significantly reduce *Pneumocystis* β-glucan induced inflammatory signaling in vitro and in vivo, suggesting that BRD5529 could be a promising PCP treatment add on agent ^2,3^.

In this study we gave mice BRD5529 at 1mg/kg or vehicle control IP daily for 7 days starting at day 42 of the PCP mouse model. Analysis of serum chemistry results suggests that BRD5529 displays similar panel parameters as compared to the vehicle control alone. One noted significant difference was that in the blood urea nitrogen (BUN) levels between the two cohorts. Treatment with BRD5529 in the PCP model significantly reduced the BUN levels as compared to the vehicle control group. Interestingly, others have reported that increased BUN levels were factors of poor prognosis in PJP patients^8^. Therefore, in addition to possible reductions in proinflammatory response via CARD9 inhibition by BRD5529, other benefits of BRD5529 might be worth exploring in PJP.

## 5. Conclusions

Our preliminary findings suggest the use of BRD5529 in the PCP mouse model as measured by blood toxicology seems like a viable agent to further explore in reducing PCP-induced proinflammatory response in the host.

